# Cancer-associated fibroblast ADAM17 mediates a feed-forward loop promoting cancer cell migration

**DOI:** 10.1101/2023.05.14.540677

**Authors:** Maria L Perciato, Simon A Whawell, Daniel W Lambert

## Abstract

One of the ways in which cancer associated fibroblasts (CAF) influence the tumour-microenvironment (TME) is by releasing soluble factors to elicit responses in nearby cells. These factors may be released by modification of the extracellular matrix (ECM), secretion from intracellular compartments, or proteolytic cleavage from the cell surface. A major mediator of proteolytic processing of cell surface proteins is the ‘a disintegrin and metalloproteinase’ (ADAM) family, commonly ADAM17. The role of ADAM17 in CAF, however, remains largely unknown.

Here, we show that expression of ADAM17 was induced in normal oral fibroblasts (NOF) and CAF by exposure to oral cancer cell-derived conditioned medium and depletion of ADAM17 reduced the ability of CAF to promote cancer cell migration by negatively regulating cancer cell-associated N-cadherin. Proteomic analysis of ADAM17-depleted CAF revealed changes in the expression of pro-tumorigenic proteins, including fibroblast growth factor 2 (FGF2). Inhibition of FGF2/FGFR1 signalling abrogated the pro-migratory effects of CAF by reducing cancer cell-associated N-cadherin, an effect rescued by addition of recombinant FGF2.

Taken together, these results indicate a novel molecular mechanism underpinning cancer cell migration in which tumour-derived factors induce ADAM17 expression in CAF, thus initiating a feed-forward loop wherein CAF release FGF2 to stimulate N-cadherin-dependent cancer cell migration.

## Introduction

Cancer associated fibroblasts (CAF) play a key role in tumour growth, metastasis and resistance to therapy in a wide range of solid tumours, including oral cancer (1). In the most common form of oral cancer, oral squamous cell carcinoma, the abundance of CAF is the single biggest determinant of prognosis (2), highlighting their utility as a source of biomarkers and, potentially, as therapeutic targets. Although a wide range of phenotypic alterations have been reported in CAF (1), the mechanisms underlying their aetiology and their effects on cancer cells remain to be fully elucidated, hampering efforts to capitalise on their therapeutic potential.

One mechanism by which CAF promote tumour progression is by engaging in cross-talk with cancer cells through the release of soluble bioactive factors (3,4). This communication is bi-directional, with cancer cells releasing factors able to influence CAF phenotype, and vice versa (5,6). Several cellular processes contribute to the generation of soluble factors implicated in CAF:cancer cell interactions, including proteolytic processing of membrane-bound proteins to release physiologically active ectodomains able to bind receptors on recipient cells. One of the major mediators of proteolytic processing is the <u>a</u> <u>d</u>isintegrin <u>a</u>nd <u>m</u>etalloproteinase (ADAM) family of enzymes (7,8).

ADAM17 is the most widely expressed member of the ADAM family and is reported to be differentially expressed in tumours and contribute to malignant progression (9–15). ADAM17 was first identified as a proteinase responsible for generating the soluble, biologically active, form of TNF-□, and is hence often referred to as TNF-□-converting enzyme (TACE) (16). Since its discovery, other cytokines and chemokines, and their receptors, have been identified as ADAM17 substrates (17). As such, ADAM17 plays a key role in immune responses (9) but its physiological roles are much broader than this owing to its ability to cleave a plethora of growth factors (18,19), clotting factors (20,21), and even viral receptors (for example, ACE2, the SARS-CoV2 receptor (22)). In cancer cells, ADAM17 is known to cleave pro-tumorigenic factors such as heparin-bound EGF (HB-EGF), TGF-α and amphiregulin, which when cleaved from the cell surface are able to bind to receptors on recipient cells and provoke both physiological and pathophysiological responses (23–25). ADAM17-mediated substrate cleavage also generates intracellular protein fragments, such as the notch intracellular C-terminal domain (ICD), that provoke nuclear responses resulting in gene expression changes in the host cell. This bi-functional activity of ADAM17, influencing the behaviour not just of the recipient cell but also the host cell, potentiates the effects of ADAM17 in cell-cell communication.

The vast majority of studies to date have concentrated on the role of ADAM17 in cancer cells (10), with little attention given to its function in CAF. Here, we hypothesised that ADAM17 contributes to the pro-tumorigenic effects of CAF. We provide evidence that cancer cell-derived factors induce ADAM17 expression and catalytic activity in CAF. Depletion of fibroblastic ADAM17 abrogated CAF-mediated cancer cell migration and was associated with a reduction in the elevated expression of cancer cell-associated N-cadherin. Among proteins displaying altered expression in ADAM17-depleted fibroblasts, FGF2 demonstrated an ability to promote cancer cell migration, implying the existence of a pro-tumorigenic ADAM17/FGF2/N-cadherin positive feedback loop between CAF and cancer cells.

## Methods

### Cell culture

Primary normal oral fibroblasts (NOF) were isolated from human gingival or buccal tissue, as previously described (26) (Sheffield Research Ethics Committee reference number 09/H1308/66) and primary human cancer associated fibroblasts (CAF) from OSCC resection specimens (27) from Sheffield Teaching Hospitals NHS Foundation Trust (Sheffield Research Ethics Committee reference number 13/NS/0120, STH17021).

Human oral squamous cell carcinoma (OSCC)-derived cell line H357 and H376 were previously established from a SCC of the tongue excised from a 74 year-old Caucasian male patient (28) and from a SCC of the floor of the mouth excised from a 40 year-old patient (28) respectively. Both cell lines were retrieved from the department biorepository and authenticated by NorthGene by the Short Tandem Repeat (STR) DNA profiling analysis based on the Deutsche Sammlung von Mikroorganismen und Zellkulturen (DSMZ) database (H357 matching percentage of 97%, H376 matching percentage of 93%). STR profiles are included in Supplementary Table 1.

Routine screening for mycoplasma testing was performed for all cell lines and cultures and only those with negative results were used for experimentation.

All cells were cultured in DMEM (D6546-500ML, Sigma-Aldrich) supplemented with 10% fetal bovine serum (FBS) (10270-106, Gibco by Life Technologies), 2 mM L-glutamine (G7513-100ML, Sigma-Aldrich) and penicillin and streptomycin (100 IU/ml and 100 μg/ml, respectively) (P0781-100ML, Sigma-Aldrich). Cells were grown at 37□°C in a humidified atmosphere with 5% CO_2_.

### Generation of the conditioned medium

Cells, under the different experimental conditions required, were incubated for 48 h in complete culture medium. After incubation, the conditioned medium (CM) was harvested, centrifuged at 1000 x g for 5 minutes and filtered through a 0.22□μm pore filter. The CM was then either used or stored at −20 °C.

### Treatment of NOF with the recombinant human TGF-β1

NOF (2 x 10^5^ cells) were seeded in 6 well plates and incubated overnight, to allow cell attachment. The following day spent medium was discarded and replaced with complete culture medium supplemented with recombinant human TGF-β1 (0.25, 0.5, 0.1, 2.5 and 5 ng/ml; 240-B-002/CF, R&DSystems, UK) for 48h at 37° C in a humidified atmosphere with 5% CO_2_.

### Transfection of siRNA

NOF and CAF seeded into 6-well plates were transfected at ∼60-70% confluence by diluting siRNAs in serum free medium in the presence of Oligofectamine 2000 transfection reagent (Invitrogen, 12252011) following the manufacturer’s recommendations. Cells were incubated for 48 h at 37□°C in a humidified atmosphere with 5% CO_2_.

All siRNAs were purchased from ThermoFisher Scientific. Details are shown in Supplementary Table 2.

### N-cadherin inhibition via ADH-1

H357 and H376 cells were seeded in 6 well plates and when at ∼80 % confluence, cells were treated either with 0.5 % DMSO (as control) or 0.2 mg/ml of ADH-1 (HY-13541, Generon) for 24 hours. The cells were kept at 37° C in a humidified atmosphere with 5% CO_2_.

### FGFR signalling pathway assessment in cancer cells

H357 and H376 cells at ∼80 % confluence were treated either with distilled water (as control) or with 20 ng/ml of Human Recombinant FGF2 (PHG0026, ThermoFisher Scientific) for 48 h. After 48 h, when cells were 90% confluent, they were treated with 0.5 % DMSO (as control) or with the FGFR inhibitor SSR128129E at three different concentrations (2.5, 5 and 10 μg/ml) (5334340001, Sigma-Aldrich) for 24 h. Cells were kept at 37° C in a humidified atmosphere with 5% CO_2_.

### EdU incorporation assay

The 5-ethynyl-2’-deoxyuridine (EdU) incorporation assay was performed using the EdU Proliferation Assay Kit (iFluor 488) (ab219801, Abcam). Cells were seeded onto coverslips and incubated with 10 μM EdU for 4 h at 37°C in a humidified atmosphere with 5% CO_2_. Afterwards, medium was discarded and cells were fixed with 200 μl of 1 x Fixative Solution for 15 minutes at RT. Cells were washed twice with 200 μl of Wash Buffer and permeabilised with 200 μl of Permeabilisation Buffer for 20 minutes at RT. Then, cells were washed twice with 200 μl of Wash Buffer and incubated with 100 μl of Reaction Mix for 30 minutes at RT protected from light. Afterwards, cells were washed twice with Wash Buffer and once with 200 μl of PBS. The coverslips were mounted onto microscope slides using the Prolong Gold Antifade Mountant with DAPI (P36390, ThermoFisher Scientific) and viewed using a Zeiss Axioplan 2 fluorescence light microscope, at 20-40x, setting the filter for Excitement/Emission between 491 and 520nm. Images were acquired using Proplus 7.0.1 image software.

### Wound healing assay

H357 and H376 cells were seeded into 6-well plates and cultured until ∼80-90% confluent. The following day, the culture medium was discarded and replaced with the desired conditioned medium or treatment. Cells were incubated for 48 h and then, using a 200 μl pipette tip inclined at around 30 degrees, a straight scratch was made per well. The cell culture medium was aspirated and cells were rinsed twice to ensure complete removal of cellular debris and detached cells. Afterwards, cells were incubated with the fresh culture medium and a first image was acquired, using the Olympus CKX41 inverted microscope, and labelled as time 0. Further images were acquired after 6 and 24 h and analysed by Fiji using the freehand tool to mark the areas of each sample.

### Transwell migration assay

Prior to the transwell migration or invasion assays, OSCC cell lines were seeded into 6 well plates and incubated with conditioned medium derived from NOF/CAF or with fresh DMEM as control. After 48 hours, when the cells were 70-90 % confluent, cells were trypsinised and counted for the assay. In a 24 well plate, 1 ml of complete DMEM was added per well and a transwell insert (662638, Greiner BIO-ONE) was placed into each well. Next, 25 x 10^^3^ cells/200 µl/insert were seeded in serum-free DMEM. The cells were then incubated for 24-48 hours at 37 °C in a humidified atmosphere with 5 % CO_2_. At the end of the incubation, the medium from each insert was discarded and the inserts were washed quickly in PBS 1X by immersion. Next, each transwell was fixed in pure methanol for 2 minutes at room temperature. The cells adhering to the insert from the side facing the lid of the plate were gently removed using a cotton swab. Subsequently, the inserts were incubated with 1% crystal violet in 0.1% acetic acid for 20 minutes at room temperature. Next, the crystal violet was removed and the inserts were washed 3 times in tap water and let dry at room temperature. The next day, each insert was incubated with 1% acetic acid for 20 minutes at room temperature on a shaker. The optical density was then measured with a TECAN plate reader (Infinite® 200 PRO), wavelength set at 590 nm, and the migration rate quantified by normalising the absorbance to the control.

### Western blotting

Cells were lysed in 50 μl of RIPA lysis buffer (sc-24948A, ChemCruz, Santa Cruz Biotechnology) supplemented with both protease and phosphatase inhibitors (04693159001, Roche).

Total protein was quantified using the Pierce bicinchoninic protein assay (BCA) kit in accordance with manufacturer’s instructions (23225, ThermoFisher Scientific). Following protein quantification, 20□µg of sample was loaded onto 4–12% SDS-polyacrylamide gels and proteins were resolved by electrophoresis in SDS-Tris-glycine buffer, at 80-100V. Proteins were then transferred onto a nitrocellulose blotting membrane (10600003, GE Healthcare Life Science).

Membranes were incubated with either 5% bovine serum albumin (BSA) (A4503-100G, Sigma Aldrich) or 5% skimmed milk (84615.0500, VWR Prolabo Chemicals) in Tris- or phosphate-buffered saline with 0.05% Tween 20 (TBS-T and PBS-T respectively), for 1 h at RT on a shaker, and then incubated with a primary antibody at 4°C overnight.

The following day, each membrane was probed with the appropriate horseradish peroxidase-conjugated secondary antibody (anti-mouse dilution 1:5,000; GTX213112-01, Gene Tex; anti-rabbit dilution 1:3000; 7074S, Cell Signaling Technology), and incubated for 1□h at room temperature. Chemiluminescent signals were visualised, after incubating each membrane with the enhanced chemo-luminescent (ECL) Clarity Western ECL substrates (1705060, BioRad) for 1-2 minutes at RT, either using a digital developer (C-Digit, LI-COR) or by exposing it to an X-ray film (Thermo Scientific) in the dark and developing using a Compact X4 Developer (Xograph Imaging Systems). All original blots are provided as source data. Details about the primary and secondary antibodies are included in Supplementary Table 3.

### RNA isolation and RT-qPCR

RNA was isolated from cell lysates using the Monarch Total RNA Miniprep Kit (T2010S, New England BioLabs) according to the manufacturer’s recommendations. Then, the RNA was quantified using a Nanodrop 1000 spectrophotometer (Thermo Scientific) and reverse-transcribed using the High Capacity cDNA Reverse Transcription Kit (Applied Biosystems, USA), according to the manufacturer’s instructions. Quantitative PCR was performed either using SYBR green (PB20.15-05, qPCRBIO) or Taqman (qPCRBIO Probe Blue Mix Lo-ROX; PB20.25-05, qPCRBIO) and using the Rotor-Gene Q PCR system (Qiagen, Germany).

Custom primers were purchased from Sigma Aldrich and Taqman primers from ThermoFisher Scientific; sequences are available in Supplementary Table 4. Relative quantitation was performed using the ΔΔC_t_ method.

### ELISA

Supernatants were harvested and centrifuged at 1000 x g to remove cell debris and dead cells and filtered through a 0.22□μm pore filter. The sandwich ELISA assay performed was against HB-EGF (catalogue number DY259B) and all the reagents used throughout the assay were prepared according to the manufacturer’s recommendations.

### Mass spectrometry

CAF were seeded in 6 well plates according to the planned treatments (scramble siRNA and ADAM17 siRNA). For each cell type, there were 3 biological replicates. Samples were harvested by mechanically scraping cells from each well in 1 ml of culture medium. The cells were centrifuged at 1000 x g for 5 min and, after discarding the supernatant, the pellet was washed 3 times in 500 µl of ice-cold PBS. After discarding the PBS, pellets were stored at −80°C until sample submission to the Mass-Spectrophotometry Facility for a Label-Free Quantification (LFQ) Mass-Spec analysis.

The raw data were then processed using the MaxQuant quantitative proteomics software package (29). The resulting text file (renamed protein_groups.txt), containing information about the protein identification and intensities, was used as one of the two input files (the second file was a text file containing the experimental set up and was renamed experimental.txt) required for the statistical analysis performed via LFQ Analyst, a web-based and free platform designed for the analysis of label-free quantitative experiments processed with MaxQuant (30). The settings used for the statistical analysis were adjusted to a p-value cut-off equal to 0.05, Log2 fold change cut-off equal to 1, imputation type bpca, type of FDR correction Benjamini Hochberg.

The two lists of proteins that originated from the two independent mass spectrometry analyses were, regardless of the p-value, compared and screened to identify the proteins whose expression was altered in the same conditions.

### Statistical analysis and reproducibility

Statistical analyses were performed using Prism software (Graphpad Software v.8 and v.9.4). Data were reported as a mean and standard deviation. When more than two groups were compared and only one factor was considered, the One-Way Analysis Of Variance (one-way ANOVA) was applied. Unpaired t-test was used to test differences between two means. A P-value of less than 0.05 was considered significant.

All experiments were performed three or more times independently and under similar conditions, except experiments shown in figures 3G-H, 5, S3E-F and S4 which were performed only twice.

## Results

### OSCC cell-derived factors upregulate ADAM17 expression in fibroblasts and promote differentiation to a CAF-like phenotype

In order to examine the effect of cancer cell-derived factors on ADAM17 expression in fibroblasts, we established a simple indirect co-culture system in which conditioned media was harvested from two OSCC-derived cell lines, the non-metastatic H357 and the metastatic H376 and applied to normal oral fibroblasts (NOF) and cancer-associated fibroblasts (CAF) (Figure 1A). Western blot analyses of NOF and CAF-derived cell lysates revealed a significant upregulation of ADAM17 protein expression in response to H357 (2.3±0.64-fold for NOF, 2.4±0.9-fold for CAF) and H376 (3.4+/-1.17-fold for NOF, 3.47+/− 1.01-fold for CAF) (Figure 1B). No significant differences in the magnitude of upregulation of ADAM17 at the protein level were observed between NOF and CAF in response to conditioned media from either cancer cell line. A similar response was observed at the transcript level, with NOF showing a 2.5+/-0.1-fold and 1.5+/-0.07-fold increase in ADAM17 mRNA expression, and CAF a 3.7+/-0.14-fold and a 1.9+/-0.17-fold increase in response to H357 and H376-derived factors, respectively (Figure 1C). At the transcript level, a significantly greater increase in ADAM17 was observed in CAF than in NOF. This increase in expression was mirrored by catalytic function, with exposure to both H357 and H376-derived conditioned media significantly increasing the shedding of a heterologously expressed ADAM17 substrate conjugated to an extracellular alkaline phosphatase tag (Figure 1D). In the case of ADAM17 catalytic activity, the increase in CAF in response to conditioned media from both cell lines was significantly greater than in NOF, although we cannot exclude the possibility that construct expression was higher in CAF so this result should be interpreted with caution. The observed increase in ADAM17 in response to OSCC-derived factors was associated with a significant increase in the expression of markers of the CAF phenotype, collagen I and FAP, at the level of protein (Figure 1E), and collagen I at transcript level (Figure 1F). These differences were more pronounced in NOF than CAF, particularly at the protein level. No change was observed in the expression of □-SMA at either protein or transcript level.

**Figure 1:**
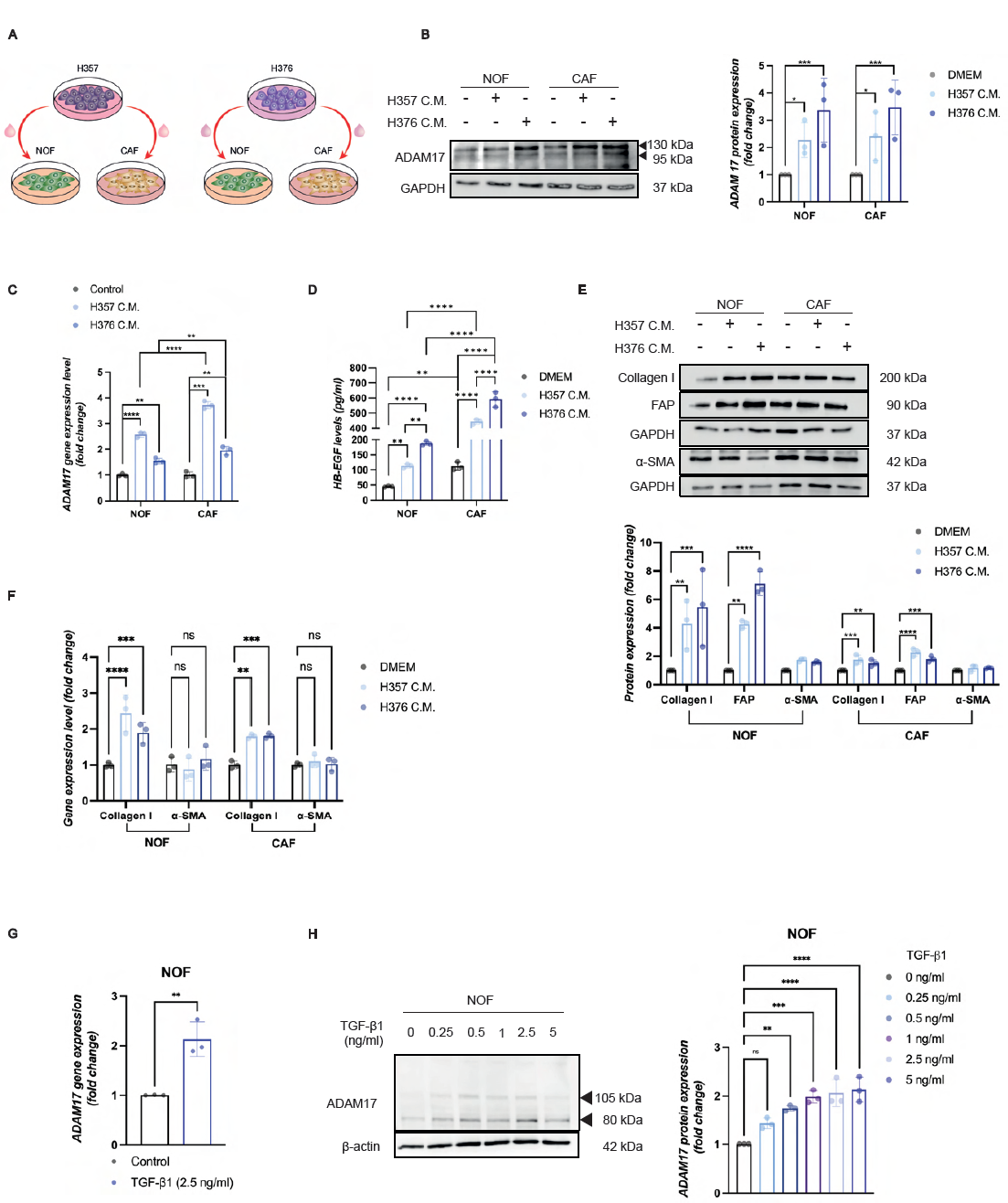
Cancer cell-derived conditioned medium triggers ADAM17 upregulation in fibroblasts and the acquisition of a CAF-like phenotype. A. Schematic representation of the experimental set up. B. Western blot showing ADAM17 expression in NOF and CAF after incubation with or without the conditioned medium derived from H357 and H376 cells and its quantification by densitometry. Representative of three independent experiments. One-way ANOVA (*P ≤ 0.05, **P ≤ 0.01, ***P ≤ 0.001, ****P ≤ 0.0001). C. RT-qPCR showing ADAM17 expression in NOF and CAF after incubation with or without the conditioned medium derived from H357 and H376 cells. Each dot represents a biological replicate from three independent experiments (*n*□=□3). One way ANOVA (*P ≤ 0.05, **P ≤ 0.01, ***P ≤ 0.001, ****P ≤ 0.0001). D. ELISA showing HB-EGF levels in the supernatant derived from NOF and CAF after incubation with or without the conditioned medium derived from H357 and H376 cells. Representative of three independent experiments. One way ANOVA (*P ≤ 0.05, **P ≤ 0.01, ***P ≤ 0.001, ****P ≤ 0.0001). E. Western blot showing collagen I, FAP and α-SMA expression in NOF and CAF after incubation with or without the conditioned medium derived from H357 and H376 cells and its quantification by densitometry. Representative of three independent experiments. One way ANOVA (*P ≤ 0.05, **P ≤ 0.01, ***P ≤ 0.001, ****P ≤ 0.0001). F. RT-qPCR showing collagen I and α-SMA expression in NOF and CAF after incubation with or without the conditioned medium derived from H357 and H376 cells. Each dot represents a biological replicate from three independent experiments (*n*□=□3). One way ANOVA (*P ≤ 0.05, **P ≤ 0.01, ***P ≤ 0.001, ****P ≤ 0.0001). G. RT-qPCR showing ADAM17 expression in NOF upon incubation with recombinant TGF-β1 (2.5 ng/ml). Each dot represents a biological replicate from three independent experiments (*n*□=□3). Unpaired t test (*P ≤ 0.05, **P ≤ 0.01, ***P ≤ 0.001, ****P ≤ 0.0001). H. Western blot showing ADAM17 expression in NOF upon incubation with recombinant TGF-β1 at different concentrations and its quantification by densitometry. Representative of three independent experiments. One way ANOVA (*P ≤ 0.05, **P ≤ 0.01, ***P ≤ 0.001, ****P ≤ 0.0001). Data are mean ± s.d. Results are obtained by normalisation to untreated samples (DMEM or control).

The association between changes in ADAM17 expression and the acquisition of CAF-like traits was further explored by assessing the effect of TGF-β1, a growth factor widely employed *in vitro* to provoke differentiation of normal fibroblasts to a CAF-like phenotype (31). TGF-treatment of NOF resulted in a dose-responsive increase in ADAM17 protein expression (maximal at 2±0.3-fold, at 2.5 ng/ml concentration), and a corresponding increase in transcript levels (Figure 1G, H). This increase in ADAM17 expression was associated with elevated levels of some CAF markers including collagen I and □-SMA, and activation of SMAD3 phosphorylation (Figure S1B, C).

### Depletion of ADAM17 inhibits the CAF phenotype

Having established that elevated ADAM17 expression is associated with the development of a CAF phenotype, we next sought to examine the impact of reducing ADAM17 expression in CAF. Depletion of ADAM17 in CAF (henceforth referred to as CAF^A17low^) was successfully achieved using siRNA (3±0.12-fold reduction in protein, 2.9+0.05-fold reduction in transcript) (Figure 2A-C). Having previously observed an increase in CAF-like markers concomitantly with the up-regulation of ADAM17 levels and catalytic activity, we assessed whether silencing ADAM17 in CAF could affect the expression of CAF-associated markers. Western blot analysis showed a significant decrease of FAP protein levels (1.3±0.05-fold, 1.9±0.49-fold and 2.2±0.16-fold in CAF^A17low^ compared to their respective controls), and the transcript levels of □SMA, vimentin and collagen I also decreased by 1.3± 0.03-fold, 1.25±0.06-fold and 1.4±0.03-fold respectively, in CAF^A17low^ compared to control (Figure 2D, E). Similar trends were observed in NOF, although the reduction of CAF-associated markers was less marked and, in some cases, not significant (Fig S2). At the protein level only FAP decreased, by 2±0.2-fold in NOF^A17low^, previously exposed to the CM derived from cancer cells (cNOF), whereas, at transcriptional level, □SMA and collagen I, but not vimentin, decreased by 1.3±0.01-fold and 1.3±0.04-fold (Figure S2 A-C).

**Figure 2:**
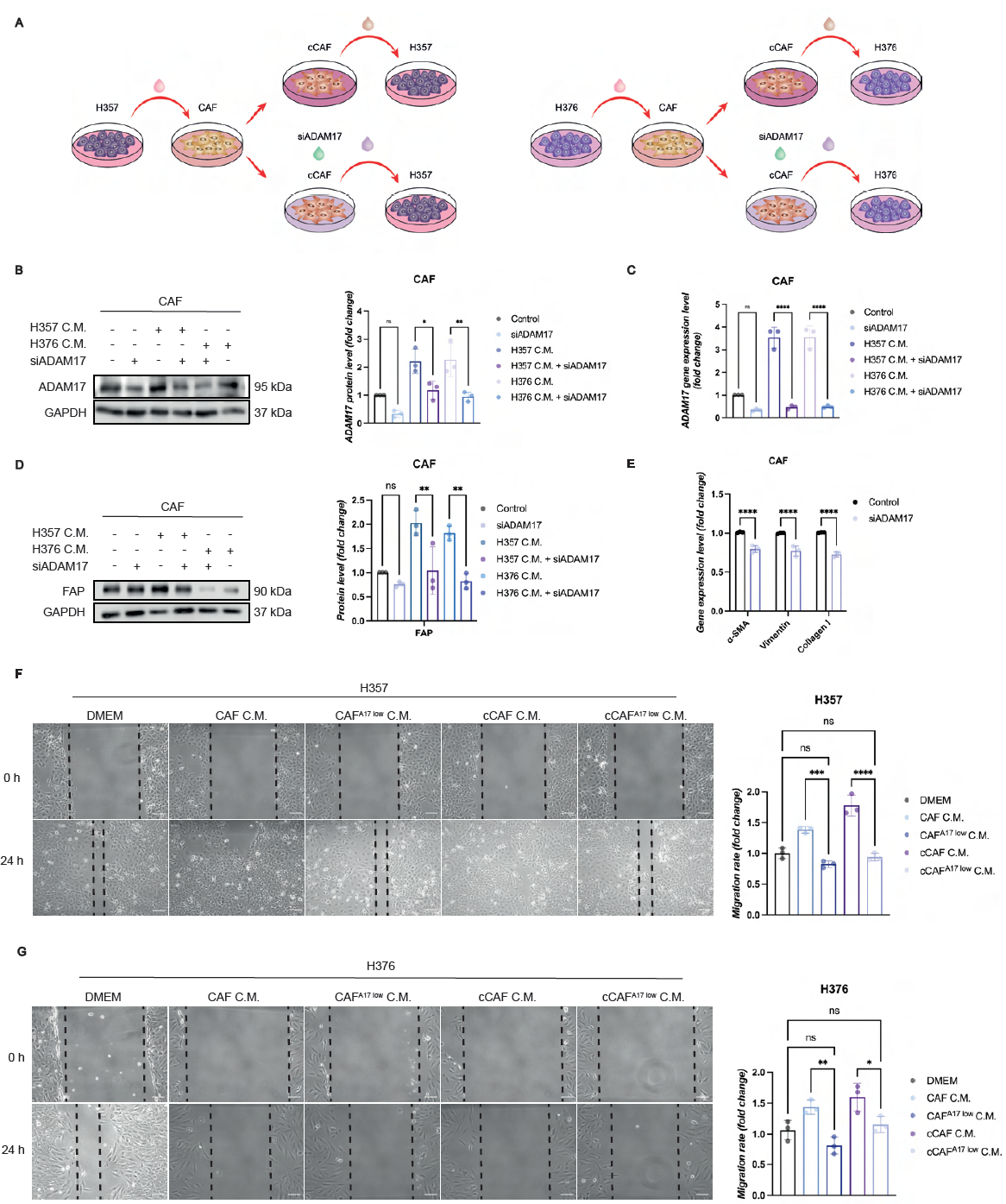
ADAM17 depletion in CAF restrains CAF properties. **A.** Schematic representation of the experimental set up. **B.** Western blot showing ADAM17 expression in CAF depleted of ADAM17 by siRNA after incubation with or without the conditioned medium derived from H357 and H376 cells and its quantification by densitometry. Representative of three independent experiments. One-way ANOVA (*P ≤ 0.05, **P ≤ 0.01, ***P ≤ 0.001, ****P ≤ 0.0001). **C.** RT-qPCR showing ADAM17 expression in CAF depleted of ADAM17 by siRNA after incubation with or without the conditioned medium derived from H357 and H376. Each dot represents a biological replicate from three independent experiments (*n*□=□3). One way ANOVA (*P ≤ 0.05, **P ≤ 0.01, ***P ≤ 0.001, ****P ≤ 0.0001). **D.** Western blot showing FAP expression in CAF depleted of ADAM17 by siRNA after incubation with or without the conditioned medium derived from H357 and H376 cells and its quantification by densitometry. Representative of three independent experiments. One-way ANOVA (*P ≤ 0.05, **P ≤ 0.01, ***P ≤ 0.001, ****P ≤ 0.0001). **E.** RT-qPCR showing collagen I, vimentin and α-SMA expression in CAF upon ADAM17 depletion by siRNA. Each dot represents a biological replicate from three independent experiments (*n*□=□3). Unpaired t test (*P ≤ 0.05, **P ≤ 0.01, ***P ≤ 0.001, ****P ≤ 0.0001). **F.** Wound healing assay showing the migration potential of H357 cells incubated with or without the conditioned medium derived from CAF and cCAF depleted or not of ADAM17. Representative images from three independent experiments. Scale bars, 200 μm. One-way ANOVA (*P ≤ 0.05, **P ≤ 0.01, ***P ≤ 0.001, ****P ≤ 0.0001). **G.** Wound healing assay showing the migration potential of H376 cells incubated with or without the conditioned medium derived from CAF and cCAF depleted or not of ADMA17. Representative images from three independent experiments. Scale bars, 200 μm. One-way ANOVA (*P ≤ 0.05, **P ≤ 0.01, ***P ≤ 0.001, ****P ≤ 0.0001). Data are mean ± s.d. Results are obtained by normalisation to untreated samples (DMEM).

Next, to evaluate whether ADAM17 depletion in CAF and NOF could have any impact on modulating cancer cell motility, a wound healing assay was performed using both H357 and H376 upon treatment with the CM derived from CAF^A17low^ and NOF^A17low^. The data showed that the migration rate of both cancer cell lines was robustly and significantly restricted when incubated with the CM derived from ADAM17 deficient CAF and NOF, and from ADAM17 deficient CAF and NOF previously exposed to cancer cell-derived CM (cCAF and cNOF respectively). Indeed, compared to control, the migration rate in H357 decreased by 1.67±0.05-fold and 1.85±0.06-fold when treated with the CM derived from CAF^A17low^ and cCAF^A17low^ respectively and in H376 it decreased by 1.8±0.14-fold and 1.4±0.13-fold (Figure 2F, G). Likewise, the motility in H376 decreased by 2.3±0.02-fold and 2±0.13-fold when treated with the CM derived from NOF^A17low^ and cNOF^A17low^ respectively (Figure S2E).

Consistently with these findings, H357 cells migrated less when incubated with the CM derived from NOF treated with TGF-β1 depleted of ADAM17 (henceforth eCAF^A17low^) (Figure S2H). However, eCAF^A17low^ did not display a decrease in □ SMA and collagen I levels (Figure S2F, G).

Altogether these findings indicate that targeting ADAM17 in CAF may contribute to a mitigation of CAF-associated pro-tumorigenic properties by negatively regulating markers such as FAP and by re-modulating CAF secretome.

### CAF-associated ADAM17 regulates cancer cell-associated N-cadherin levels

To determine the potential mechanism underpinning the CAF-associated ADAM17 modulation of cancer cell migration, we assessed the expression levels of four genes (Snail, Slug, Twist and N-cadherin) associated with epithelial-mesenchymal transition (EMT) and the acquisition of a motile capability by cancer cells (32). By qPCR analysis, N-cadherin was the only gene showing a significant reduction in expression in both H357 and H376 cells in response to the CM derived from CAF^A17low^ and cCAF^A17low^ (1.7±0.1-fold and 1.6±0.24-fold in H357, 1.5±0.03-fold and 1.9±0.11-fold in H376) (Figure 3A, B). By western blot the same trend was observed at protein level in both cancer cell lines under the same conditions (2.5±0.3-fold and 2.8±0.05-fold in H357, 1.9±0.07-fold and 4.5±0.3-fold) (Figure 3C, D; Figure S3A, B). Of note, although N-cadherin acquisition is generally associated with the EMT process, concomitantly with reduction of E-cadherin levels, here we did not detect any change in E-cadherin levels (Figure S3C, D).

**Figure 3:**
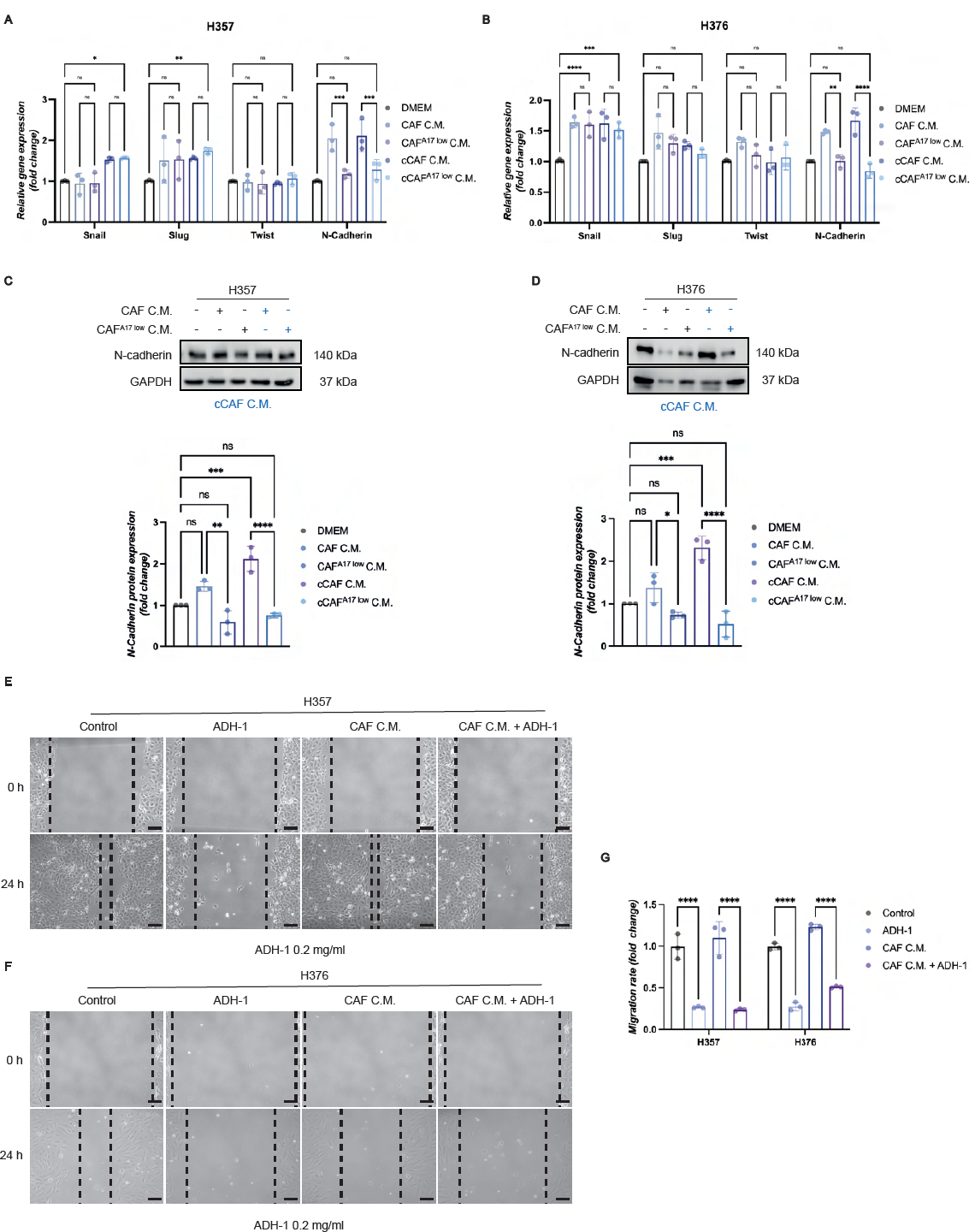
ADAM17-depleted CAF mediate N-cadherin downregulation in cancer cells. **A.** RT-qPCR showing snail, slug, twist and N-cadherin expression in H357 cells incubated with or without the conditioned medium derived from CAF and cCAF depleted of ADAM17 by siRNA. Each dot represents a biological replicate from three independent experiments (*n*□=□3). One-way ANOVA (*P ≤ 0.05, **P ≤ 0.01, ***P ≤ 0.001, ****P ≤ 0.0001). **B.** RT-qPCR showing snail, slug, twist and N-cadherin expression in H376 cells incubated with or without the conditioned medium derived from CAF and cCAF depleted of ADAM17 by siRNA. Each dot represents a biological replicate from three independent experiments (*n*□=□3). One-way ANOVA (*P ≤ 0.05, **P ≤ 0.01, ***P ≤ 0.001, ****P ≤ 0.0001). **C.** Western blot showing N-cadherin expression in H357 cells incubated with or without the conditioned medium derived from CAF and cCAF depleted of ADAM17 by siRNA, and its quantification by densitometry. Representative of three independent experiments. One-way ANOVA (*P ≤ 0.05, **P ≤ 0.01, ***P ≤ 0.001, ****P ≤ 0.0001). **D.** Western blot showing N-cadherin expression in H376 cells incubated with or without the conditioned medium derived from CAF and cCAF upon depletion of ADAM17 by siRNA, and its quantification by densitometry. Representative of three independent experiments. One-way ANOVA (*P ≤ 0.05, **P ≤ 0.01, ***P ≤ 0.001, ****P ≤ 0.0001). **E.** Wound healing assay showing the migration potential of H357 cells incubated with or without the conditioned medium derived from CAF and challenged with the N-cadherin antagonist ADH-1 (0.2 mg/ml). Representative images from two independent experiments. Scale bars, 200 μm. **F.** Wound healing assay showing the migration potential of H376 cells incubated with or without the conditioned medium derived from CAF and challenged with the N-cadherin antagonist ADH-1 (0.2 mg/ml). Representative images from two independent experiments. Scale bars, 200 μm. **G.** Quantification of **E** and **F**. One-way ANOVA (*P ≤ 0.05, **P ≤ 0.01, ***P ≤ 0.001, ****P ≤ 0.0001). Data are mean ± s.d. Results are obtained by normalisation to untreated samples (DMEM or control).

The ability of N-cadherin to modulate cancer cell migration in response to CAF-derived factors was tested by inhibiting N-cadherin in both H357 and H376 using the N-cadherin antagonist ADH-1 (33,34). As shown by a wound healing assay, cancer cells, in the absence or presence of the CM derived from CAF, upon treatment with ADH-1 displayed a robust and significant reduction in their migration (3.7±0.01-fold and 4.6±0.01-fold in H357 and 3.6±0.05 and 2.4±0.01 in H376) (Figure 3E-G). This restrained migration was independent of proliferation as no significant alteration in proliferation was observed in either cell line (Figure S3E, F).

In summary, these data point to a potential role for CAF-associated ADAM17 in mediating cancer cell migration by regulating cancer cell-associated N-cadherin levels.

### ADAM17 depletion in CAF negatively regulates cancer cell-associated N-cadherin via the FGFR signalling pathway

To gain further insights into the mechanisms behind the CAF-associated ADAM17-mediated regulation of cancer cell migration, we performed two independent mass spectrometry analyses to identify any differentially expressed proteins between CAF and CAF^A17^ ^low^. This approach identified 181 differentially expressed proteins of which 36 were downregulated in CAF^A17low^ (Figure 4A). Amongst the downregulated proteins,15 proteins clustered together as illustrated by protein-protein interaction (PPI) network functional enrichment analysis obtained via the open-source STRING (Figure 4B). This cluster comprised of ACTN4, CALD1, COL12A1, COL1A1, CSRP2, CSTB, ERLIN1, FBN1, FBN2, FGF2, HEBP2, LAMA4, NES, PDGFRA and TNXL1, together involved in the regulation of focal adhesion, amoebiasis, PI3K/AKT signalling pathway and actin cytoskeleton (Figure 4C). Of note, amongst these proteins, FGF2, NES, PDGFRA are involved in the regulation of the FGFR signalling pathway and have been reported to mutually and positively regulate each other (35,36). CAF have been reported to shape a pro-tumorigenic environment through the release of FGF ligands which then bind to and activate their cognate receptors on the surface of cancer cells (37–39). Moreover, CAF have been shown to promote cancer cell migration and invasion in colorectal cancer cells, in a cell-contact dependent manner, through CAF surface-associated FGF2 (38). Therefore, these data prompted us to further investigate the role of CAF-associated FGF2 in cancer cell migration as a downstream effector of the CAF-associated ADAM17:cancer cell-N-cadherin axis.

**Figure 4:**
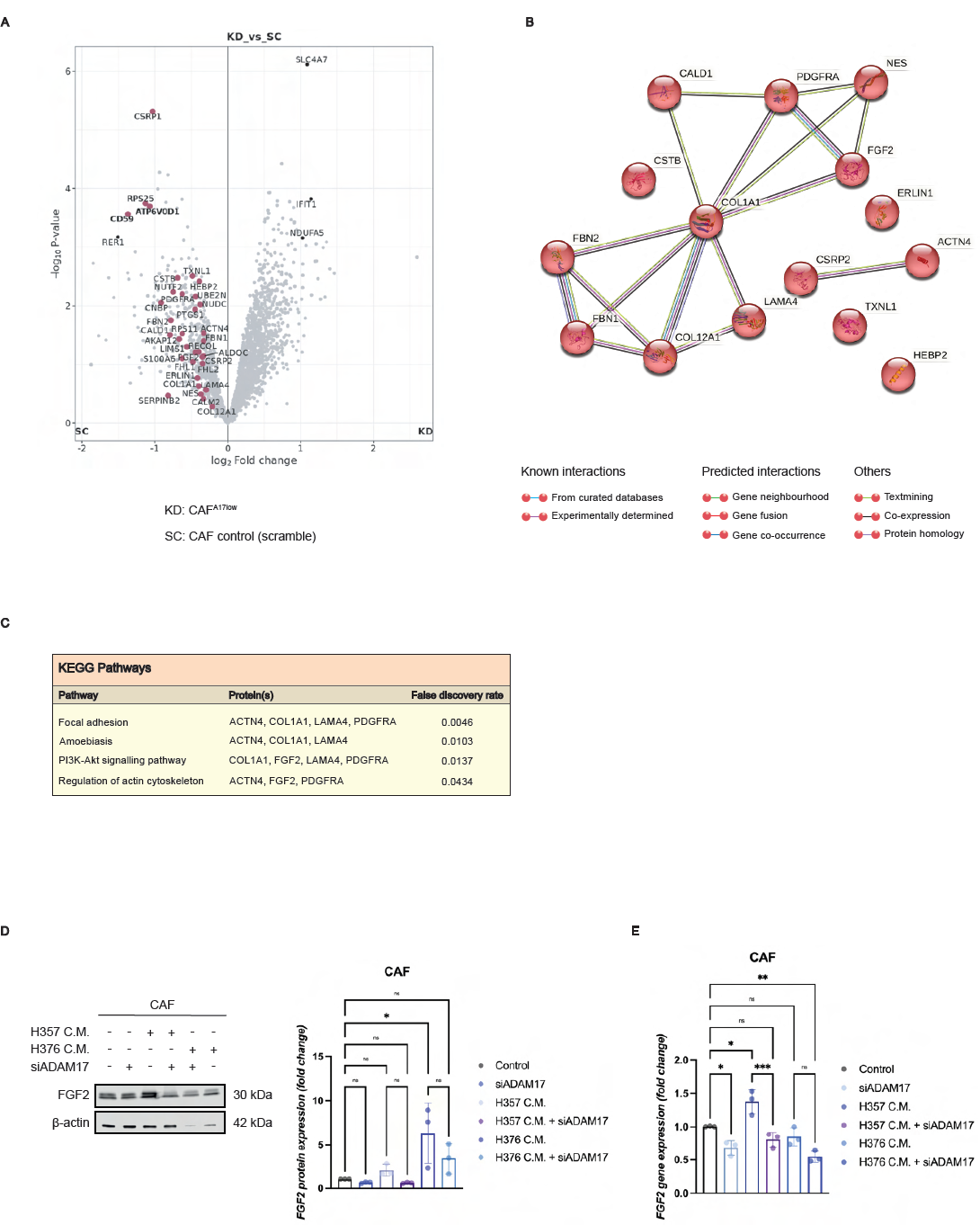
Depletion of ADAM17 in CAF downregulates FGF2. **A.** Volcano plots showing the Differentially Expressed Proteins (DEP) in CAF upon depletion of ADAM17 by siRNA. Adjusted p-value cut off, 0.05 (Benjamini-Hochberg method). Log2 fold, 1. **B.** Protein-protein interaction (PPI) network showing the proteins down-regulated upon ADAM17 depletion in CAF. **C.** KEGG analysis of DEP cluster, associated to FGF2 functions, in CAF upon depletion of ADAM17 by siRNA. **D.** Western blot showing FGF2 expression in CAF and cCAF upon depletion of ADAM17 by siRNA, and its quantification by densitometry. Representative of three independent experiments. One-way ANOVA (*P ≤ 0.05, **P ≤ 0.01, ***P ≤ 0.001, ****P ≤ 0.0001). **E.** RT-qPCR showing FGF2 expression in CAF and cCAF upon depletion of ADAM17 by siRNA. Each dot represents a biological replicate from three independent experiments (*n*□=□3).One-way ANOVA (*P ≤ 0.05, **P ≤ 0.01, ***P ≤ 0.001, ****P ≤ 0.0001). Data are mean ± s.d. Results are obtained by normalisation to untreated samples (control).

Assessment of CAF-associated FGF2 levels, by qPCR and western blot analyses, indicated its upregulation upon exposure to the cancer cell-derived CM (at transcript level with H357 CM and protein level with H376 CM) and its downregulation upon ADAM17 depletion (Figure 4D, E). In light of these findings, we next tested cancer cell migration in response to modulating FGF2. Firstly, we evaluated the impact of inhibiting cancer cell-associated FGFR with the FGFR inhibitor (FGFRi) SSR128129E at two different concentrations (5 and 10 µg/ml). As shown by wound healing assay, the migratory behaviour of both H357 and H376 cells was restrained even in the presence of the CM derived from CAF (Figure 5A, B. Likewise, N-cadherin levels showed a significant reduction in both cell lines under the same conditions (Figure 5C).

**Figure 5:**
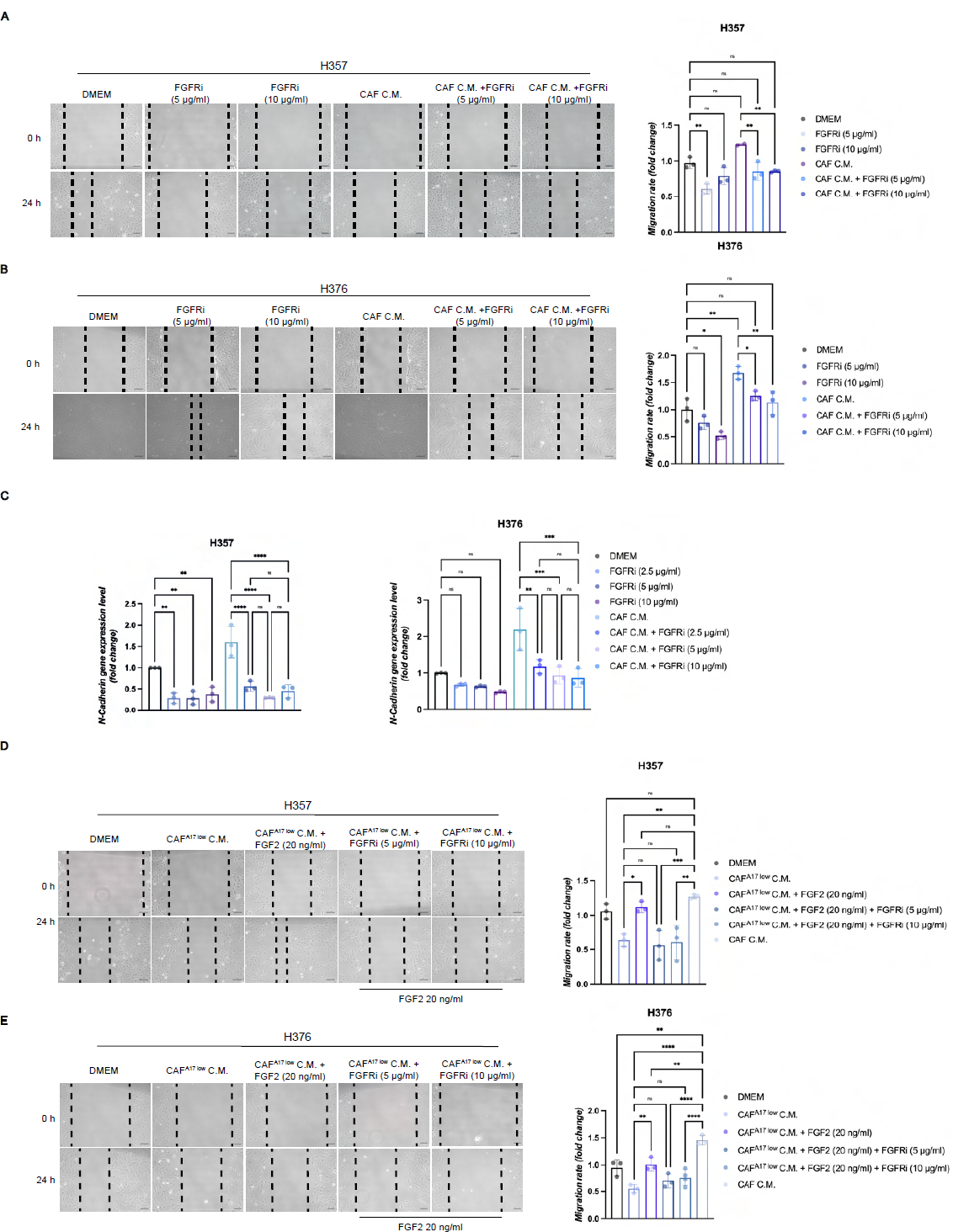
The FGF2:FGFR axis mediates cancer cell migration. **A.** Wound healing assay showing the migration potential of H357 cells incubated with or without the conditioned medium derived from CAF and challenged with the FGFR inhibitor (FGFRi) SSR128129E (5 and 10 μg/ml). Representative of two independent experiments. Scale bars, 200 μm. One-way ANOVA (*P ≤ 0.05, **P ≤ 0.01, ***P ≤ 0.001, ****P ≤ 0.0001). **B.** Wound healing assay showing the migration potential of H376 cells incubated with or without the conditioned medium derived from CAF and challenged with the FGFR inhibitor (FGFRi) SSR128129E (2.5, 5 and 10 μg/ml). Representative of two independent experiments performed in technical triplicates. Scale bars, 200 μm. One-way ANOVA (*P ≤ 0.05, **P ≤ 0.01, ***P ≤ 0.001, ****P ≤ 0.0001). **C.** RT-qPCR showing N-cadherin expression in H357 and H376 cells incubated with or without the conditioned medium derived from CAF and challenged with the FGFR inhibitor (FGFRi) SSR128129E. One-way ANOVA (*P ≤ 0.05, **P ≤ 0.01, ***P ≤ 0.001, ****P ≤ 0.0001). **D.** Wound healing assay showing the migration potential of H357 cells incubated with or without the conditioned medium derived from CAF^A17^ ^low^, then treated with the human recombinant FGF2 (20 ng/ml) and challenged with the FGFR inhibitor (FGFRi) SSR128129E (5 and 10 μg/ml). Representative of two independent experiments performed in technical triplicates. Scale bars, 200 μm. One-way ANOVA (*P ≤ 0.05, **P ≤ 0.01, ***P ≤ 0.001, ****P ≤ 0.0001). **E.** Wound healing assay showing the migration potential of H376 cells incubated with or without the conditioned medium derived from CAF^A17^ ^low^, then treated with the human recombinant FGF2 (20 ng/ml) and challenged with the FGFR inhibitor (FGFRi) SSR128129E (2.5, 5 and 10 μg/ml). Representative of two independent experiments performed in technical triplicates. Scale bars, 200 μm. One-way ANOVA (*P ≤ 0.05, **P ≤ 0.01, ***P ≤ 0.001, ****P ≤ 0.0001). Data are mean ± s.d. Results are obtained by normalisation to untreated samples (DMEM).

Next, we sought to examine the direct effects of FGF2 in mediating cancer cell migration by incubating both H357 and H376 with a human recombinant FGF2 (hrFGF2). Prior to FGF2 treatment, cancer cells were incubated with or without the CM derived from CAF and CAF^A17low^. At the end of the incubation time, cancer cells were treated with or without the FGFRi. hrFGF2 restored the motility of both H357 and H376 incubated with the CM derived from CAF^A17low^ to levels comparable to their respective controls (Figure 5D, E). Of note, the proliferative behaviour of both cancer cell lines did not change across any conditions (Figure S5).

Taken together, these data suggest that CAF-associated ADAM17 might regulate cancer cell migration by modulating the levels of CAF-associated FGF2 which, in turn, promote cancer cell migration by upregulating cancer cell-associated N-cadherin via the activation of the cancer cell-associated FGFR signalling pathway.

## Discussion

Metastasis still represents the major cause of death from cancer despite the recent advances in cancer research (40–42). Challenges in designing effective therapeutic strategies include the lack of specific targets and the limited understanding of the mechanisms underlying metastasis (40,42–44).

Moreover, large-scale genomic sequencing data have not revealed *de novo* mutations associated with the spawning of metastasis, thus raising the possibility of a cancer-cell extrinsic contribution in mediating metastasis formation (45–47).

In this regard, the role of CAF as pivotal orchestrators of cancer progression has been acknowledged and recent studies have shed light on some of the CAF-associated pro-tumorigenic functions (5,48–50). Importantly, it has been demonstrated that cancer cells and CAF engage in bi-directional communication wherein they mutually influence each other’s behaviour and fate (48,51–53). Therefore, recent approaches to tackle cancer progression focussed on targeting the factors derived from cancer cells responsible of fostering CAF formation and/or those derived from CAF (1,54–57).

In this scenario, a critical contribution is played by ADAMs such as ADAM17, known to modulate numerous biological processes by their proteolytic activity. Despite having been intensively investigated as a cancer cell-intrinsic factor, it has recently been observed to be upregulated in CAFs adjacent to the neoplastic lesions, suggesting a potential role of ADAM17 in CAF-mediated modulation of cancer cell behaviour (58–60).

Here we show that CAF-associated ADAM17 plays a critical role in cancer cell-CAF crosstalk by inducing CAFs to release pro-tumorigenic factors which render cancer cells more motile, and that cancer cells upregulate the expression of ADAM17 in CAF.

Recently, two independent studies have reported the upregulation of ADAM17 in CAFs compared with donor-matched fibroblasts, demonstrating its association with the exacerbation of cancer cell traits (59,60). However, neither of them identified the mechanisms underpinning both ADAM17 upregulation in CAF and its pro-tumorigenic implications in a CAF-dependent manner. In our study, we demonstrated that the conditioned medium derived from cancer cells induced fibroblasts to acquire a CAF-like phenotype, as shown by the increased expression of CAF-associated markers such as collagen I and FAP and, more importantly, it exacerbated CAF-like features in already reactive fibroblast by rendering them more tumour-supportive through the upregulation and stimulation of ADAM17.

Moreover, unlike Gao et al. (59) but in line with what reported by Ishimoto et al. (60), we showed that CAF-associated ADAM17 upregulation modifies the CAF-secretome by enriching it with factors which foster cancer cell migration and invasion without affecting proliferation. Of note, a novel finding in this study is that ADAM17 upregulation in CAF can be promoted by cancer cell-derived factors such as TGF-β1 and that ADAM17 depletion in CAF was able to negatively regulate the expression of CAF-associated markers (FAP) and to reduce the pro-tumorigenic functions of CAF by remodulating their secretome which rendered cancer cells less motile by downregulating N-cadherin levels.

Cancer cell migration is generally associated with the EMT programme which is mediated by several transcription factors responsible for the transition from the epithelial to a mesenchymal state by switching from E-cadherin to N-cadherin (61). In this regard, CAF have been shown to modulate the EMT program through the regulation of the EMT-associated transcription factors (62–65). Despite cancer cell-associated ADAM17 has been reported to promote EMT in diverse tumour types, the role of CAF-associated ADAM17 as EMT regulator has never been elucidated (66–68). For the first time, we demonstrated that, amongst a panel of EMT markers, N-cadherin was the only one displaying a robust and significant increase at RNA level in both H357 and H376 cells upon incubation with the CM derived from CAF, and the only one to be negatively regulated upon incubation of cancer cells with the CM derived from CAF^A17low^. Interestingly, E-cadherin levels did not fluctuate across conditions suggesting that its loss might be dispensable during the CAF-associated ADAM17-mediated migration of cancer cells.

This finding, pointing at a pivotal role of N-cadherin in cancer cell migration, was further confirmed at protein level and N-cadherin role in cancer cell migration was then validated via its inhibition using the N-cadherin antagonist ADH-1. We found that ADH-1 treatment of cancer cells inhibited their migration also when previously incubated with the CM derived from CAF.

Moreover, following proteome profiling analysis performed on CAF and CAF^A17low^, we found that, amongst the differentially expressed proteins, the protein levels of FGF2, known to be involved in the regulation of N-cadherin levels (69,70), were lower in CAF^A17low^ compared with CAF. FGF2 downregulation was subsequently confirmed by western blot and qPCR analyses.

Thus, we firstly validated the potential role of the FGF2/FGFR signalling in the paracrine mediated regulation of cancer cells by chemically targeting FGFR in cancer cells using an FGFR inhibitor and by treating cancer cells, previously incubated with the CM derived from CAF^A17low^, with the hrFGF2. Indeed, FGFR inhibition mimicked the effect of the CM derived from CAF^A17low^ by restraining cancer migration and downregulating N-cadherin RNA levels, whereas hrFGF2 was sufficient to restore the migratory potential of cancer cells previously incubated with the CM derived from CAF^A17low^.

Of note, previous studies have reported ADAM17 upregulation in response to stimulation with TGF-β1 within cancer cells (71,72), however, we demonstrated, for the first time in fibroblasts, that eCAF derived from indolent fibroblasts upon treatment with the hrTGF-β1 displayed increased RNA and protein levels of ADAM17 which were also associated with a pro-migratory secretome which rendered cancer cells more motile. Similarly to what was observed in patients-derived CAF, ADAM17 depletion in eCAF recapitulated the restriction of cancer cell migration observed with the CM derived from CAF^A17low^, thus indicating TGF-β1 as one of the potential cancer cell-associated secreted factors released by cancer cells to generate CAF expressing high levels of ADAM17.

Noteworthy, we showed for the first time, that CAF-associated, and not cancer cell-associated, ADAM17, can exert a paracrine regulation of cancer cell migration, therefore implying a cell type-dependent function of ADAM17 which is of crucial relevance for designing future therapeutic strategies.

Altogether these findings unravel for the first time one of the potential sequences of events underpinning the cancer cells:CAF crosstalk responsible for the orchestration of a pro-tumorigenic environment. We show that cancer cells, by releasing factors such as TGF-β1, promote the differentiation of CAF from indolent fibroblasts also by inducing them to upregulate ADAM17 which, in turn, leads to the generation of a tumour supportive secretome by the release of FGFR ligands such as FGF2 which eventually, upon binding to the cancer cell-associated FGFR, trigger the upregulation of N-cadherin thus promoting cancer cell migration.

Of relevance to clinical translation, we found that targeting of ADAM17 in indolent fibroblasts restrained the basal motile capability of cancer cells, suggesting that fibroblast-associated ADAM17 inhibition within the neoplastic stroma could be important to prevent primary tumours from progressing. Although this aspect remains speculation, which would need further investigation, we believe that depriving fibroblasts of ADAM17 might reduce their susceptibility to cancer cell-derived stimuli responsible for their ADAM17-mediated activation.

Importantly, by identifying some of the key signalling pathways involved in the detrimental crosstalk between cancer cells:CAF, our study paves the way for a potential combinatorial therapeutic strategy by targeting CAF-associated ADAM17/FGF2 and cancer cell-associated N-cadherin/FGFR. Even though this concept requires *in vivo* validation, its application might represent a valuable clinical strategy to further investigate.

## Acknowledgements

The authors would like to acknowledge Dr Helen Colley and Dr Amy Harding for providing primary cells, Brenka McCabe for technical support and the University of Sheffield for funding this study.

