## Supplementary figure legend for "Cancer-associated fibroblast ADAM17 mediates a feed-forward loop promoting cancer cell migration"

**Supplementary figure 1.**

**A.** Schematic representation of the experimental set up. **B.** Western blot showing collagen I and the phosphorylated and non-phosphorylated SMAD3 expression in NOF upon incubation with recombinant TGF-β1 at different concentrations and its quantification by densitometry. Representative of three independent experiments. One way ANOVA (∗P ≤ 0.05, ∗∗P ≤ 0.01, ∗∗∗P ≤ 0.001, ∗∗∗∗P ≤ 0.0001). **C.** RT-qPCR showing α-SMA and collagen I expression in NOF upon incubation with the human recombinant TGF-β1 at different concentrations. Each dot represents a biological replicate from three independent experiments (*n* = 3). One way ANOVA (∗P ≤ 0.05, ∗∗P ≤ 0.01, ∗∗∗P ≤ 0.001, ∗∗∗∗P ≤ 0.0001).

Data are mean ± s.d. Results are obtained by normalisation to untreated samples (DMEM).

**Supplementary figure 2.**

**A.** RT-qPCR showing ADAM17 expression in NOF and cNOF upon depletion of ADAM17 by siRNA. Each dot represents a biological replicate from three independent experiments (*n* = 3). One-way ANOVA (∗P ≤ 0.05, ∗∗P ≤ 0.01, ∗∗∗P ≤ 0.001, ∗∗∗∗P ≤ 0.0001). **B.** Western blot showing collagen I and FAP expression in NOF and cNOF upon depletion of ADAM17 by siRNA, and its quantification by densitometry. Representative of three independent experiments. **C.** RT-qPCR showing α-SMA, vimentin and collagen I expression in NOF upon depletion of ADAM17 by siRNA. Each dot represents a biological replicate from three independent experiments (*n* = 3). Unpaired t test (∗P ≤ 0.05, ∗∗P ≤ 0.01, ∗∗∗P ≤ 0.001, ∗∗∗∗P ≤ 0.0001). **D.** ELISA showing HB-EGF levels in the supernatant derived from NOF and CAF upon depletion of ADAM17 by siRNA. Representative of three independent experiments. One way ANOVA (∗P ≤ 0.05, ∗∗P ≤ 0.01, ∗∗∗P ≤ 0.001, ∗∗∗∗P ≤ 0.0001). **E.** Wound healing assay showing the migration potential of H376 cells incubated with or without the conditioned medium derived from NOF and cNOF depleted of ADMA17 by siRNA. Representative images from three independent experiments. Scale bars, 200 μm. One way ANOVA (∗P ≤ 0.05, ∗∗P ≤ 0.01, ∗∗∗P ≤ 0.001, ∗∗∗∗P ≤ 0.0001). **F.** Western blot showing ADAM17, α-SMA and collagen I expression in NOF after treatment with the human recombinant TGF-β1 followed by ADAM17 depletion by siRNA, and its quantification by densitometry. Representative of three independent experiments. One-way ANOVA (∗P ≤ 0.05, ∗∗P ≤ 0.01, ∗∗∗P ≤ 0.001, ∗∗∗∗P ≤ 0.0001). **G.** RT-qPCR showing α-SMA and collagen I expression in NOF after treatment with the human recombinant TGF-β1 followed by ADAM17 depletion by siRNA. Each dot represents a biological replicate from three independent experiments (*n* = 3). One-way ANOVA (∗P ≤ 0.05, ∗∗P ≤ 0.01, ∗∗∗P ≤ 0.001, ∗∗∗∗P ≤ 0.0001). **H.** Wound healing assay showing the migration potential of H357 cells incubated with or without the conditioned medium derived from NOF and eCAF. Representative of three independent experiments. One-way ANOVA (∗P ≤ 0.05, ∗∗P ≤ 0.01, ∗∗∗P ≤ 0.001, ∗∗∗∗P ≤ 0.0001).

Data are mean ± s.d. Results are obtained by normalisation to untreated samples (control).

**Supplementary figure 3.**

**A.** Western blot showing slug expression in H357 cells incubated with the condition medium from CAF depleted of ADAM17 by siRNA, and its quantification by densitometry. Representative of three independent experiments. One way ANOVA (∗P ≤ 0.05, ∗∗P ≤ 0.01, ∗∗∗P ≤ 0.001, ∗∗∗∗P ≤ 0.0001). **B.** Western blot showing slug expression in H376 cells incubated with the condition medium from CAF depleted of ADAM17 by siRNA, and its quantification by densitometry. Representative of three independent experiments. One way ANOVA (∗P ≤ 0.05, ∗∗P ≤ 0.01, ∗∗∗P ≤ 0.001, ∗∗∗∗P ≤ 0.0001). **C.** Western blot showing E-cadherin expression in H357 cells incubated with the condition medium from CAF depleted of ADAM17 by siRNA, and its quantification by densitometry. Representative of three independent experiments. One way ANOVA (∗P ≤ 0.05, ∗∗P ≤ 0.01, ∗∗∗P ≤ 0.001, ∗∗∗∗P ≤ 0.0001). **D.** RT-qPCR showing E-cadherin expression in H357 cells incubated with the condition medium from CAF depleted of ADAM17 by siRNA. Each dot represents a biological replicate from three independent experiments (*n* = 3). One way ANOVA (∗P ≤ 0.05, ∗∗P ≤ 0.01, ∗∗∗P ≤ 0.001, ∗∗∗∗P ≤ 0.0001). **E.** Immunofluorescence staining of Ki67 to determine the proliferation potential of H357 cells incubated with or without the conditioned medium derived from CAF and then challenged with the N-cadherin antagonist ADH-1 (0.2 mg/ml). Representative of two independent experiments. One-way ANOVA (∗P ≤ 0.05, ∗∗P ≤ 0.01, ∗∗∗P ≤ 0.001, ∗∗∗∗P ≤ 0.0001). Ki67 (green), DAPI (blue). Scale bars, 200 μm. **F.** Immunofluorescence staining of Ki67 to determine the proliferation potential of H376 cells incubated with or without the conditioned medium derived from CAF and then challenged with the N-cadherin antagonist ADH-1 (0.2 mg/ml). Representative of two independent experiments. One-way ANOVA (∗P ≤ 0.05, ∗∗P ≤ 0.01, ∗∗∗P ≤ 0.001, ∗∗∗∗P ≤ 0.0001). Ki67 (green), DAPI (blue). Scale bars, 200 μm.

Data are mean ± s.d. Results are obtained by normalisation to untreated samples (DMEM).

**Supplementary figure 4.**

**A.** Immunofluorescence staining of Ki67 to determine the proliferation potential of H357 cells incubated with or without the conditioned medium derived from CAF and then challenged with the FGFR inhibitor (5 μg/ml and 10 μg/ml). Representative of two independent experiments. One-way ANOVA (∗P ≤ 0.05, ∗∗P ≤ 0.01, ∗∗∗P ≤ 0.001, ∗∗∗∗P ≤ 0.0001). Ki67 (green), DAPI (blue). Scale bars, 200 μm. **B.** Immunofluorescence staining of Ki67 to determine the proliferation potential of H376 cells incubated with or without the conditioned medium derived from CAF and then challenged with the FGFR inhibitor (5 μg/ml and 10 μg/ml). Representative of two independent experiments. One-way ANOVA (∗P ≤ 0.05, ∗∗P ≤ 0.01, ∗∗∗P ≤ 0.001, ∗∗∗∗P ≤ 0.0001). Ki67 (green), DAPI (blue). Scale bars, 200 μm. **C.** Immunofluorescence staining of Ki67 to determine the proliferation potential of H357 cells incubated with or without the conditioned medium derived from CAF^A17 low^ and then challenged with the hrFGF2 (20 ng/ml) and FGFR inhibitor (5 μg/ml and 10 μg/ml). Representative of two independent experiments. One-way ANOVA (∗P ≤ 0.05, ∗∗P ≤ 0.01, ∗∗∗P ≤ 0.001, ∗∗∗∗P ≤ 0.0001). Ki67 (green), DAPI (blue). Scale bars, 200 μm. **D.** Immunofluorescence staining of Ki67 to determine the proliferation potential of H376 cells incubated with or without the conditioned medium derived from CAF^A17 low^ and then challenged with the hrFGF2 (20 ng/ml) and FGFR inhibitor (5 μg/ml and 10 μg/ml). Representative of two independent experiments. One-way ANOVA (∗P ≤ 0.05, ∗∗P ≤ 0.01, ∗∗∗P ≤ 0.001, ∗∗∗∗P ≤ 0.0001). Ki67 (green), DAPI (blue). Scale bars, 200 μm.

Data are mean ± s.d. Results are obtained by normalisation to untreated samples (DMEM).
