## Supplementary figures and images for "Cancer-associated fibroblast ADAM17 mediates a feed-forward loop promoting cancer cell migration"

### Supplementary figure 1

Supplementary Figure 1

**A**

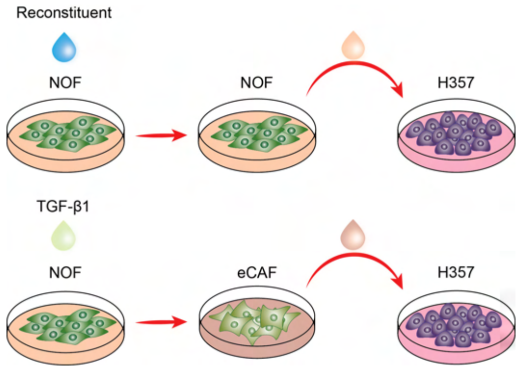


**B**

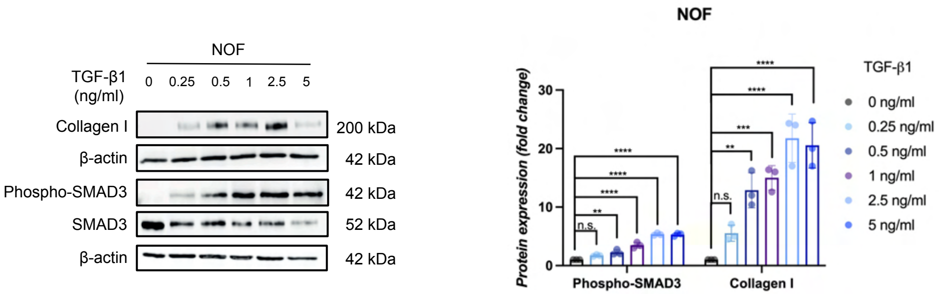


**C**

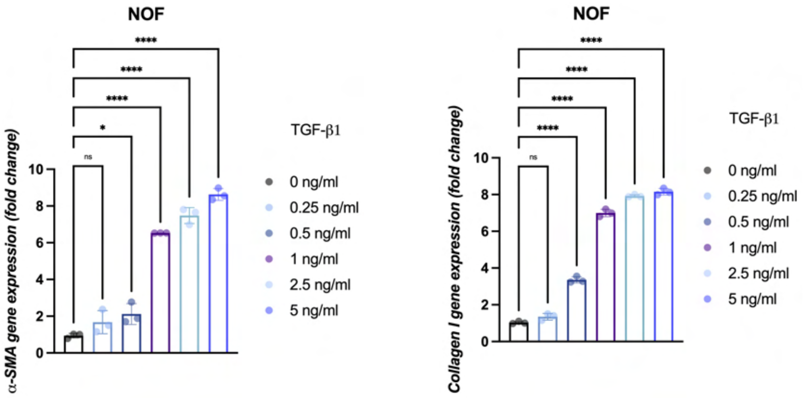

### Supplementary figure 2

#### Supplementary Figure 2

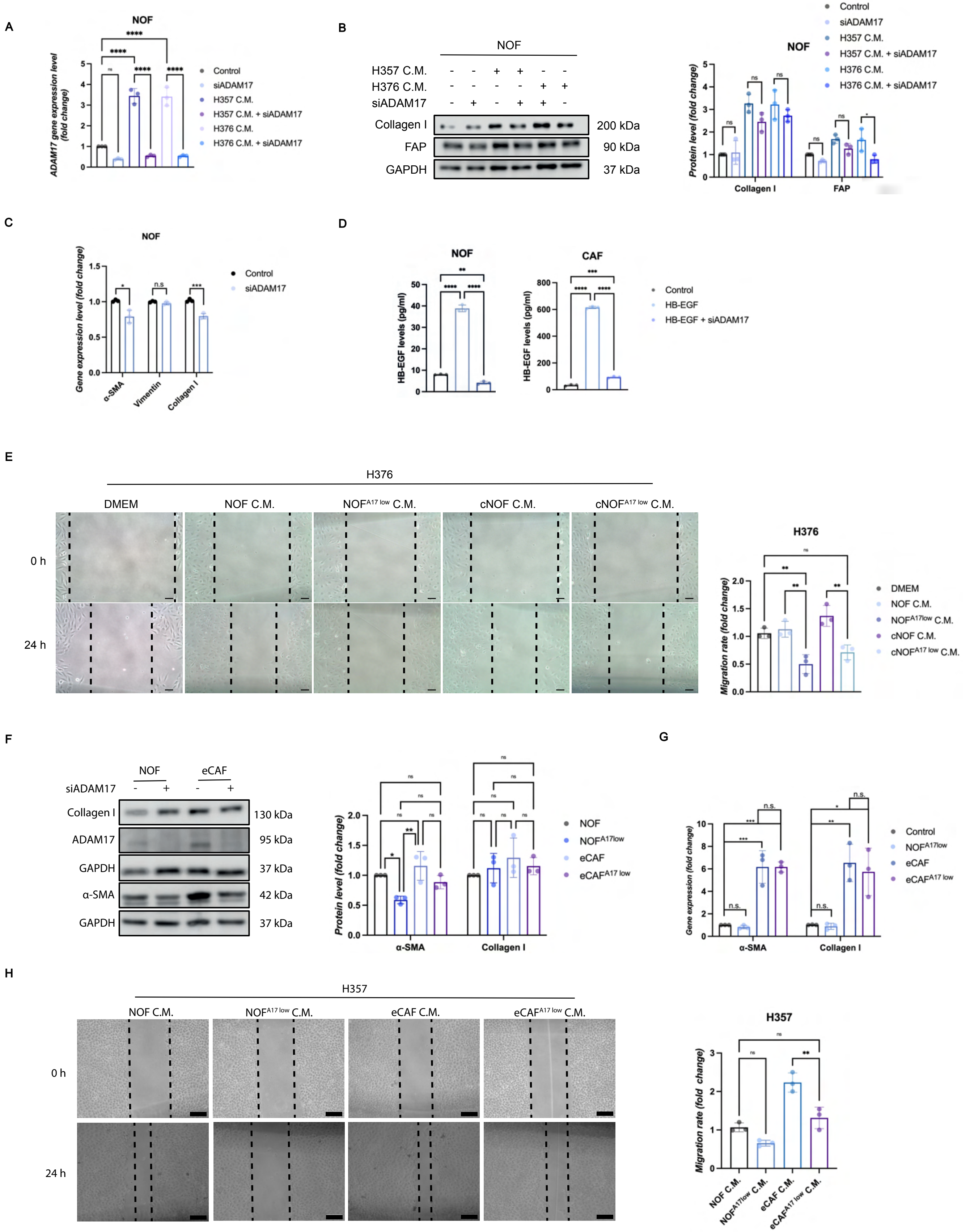

### Supplementary figure 3

#### Supplementary Figure 3

A

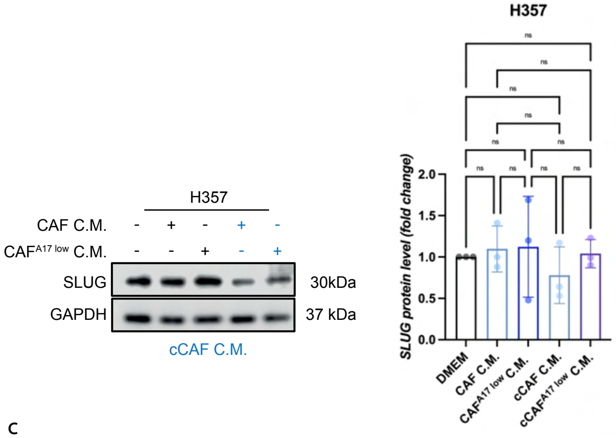


B

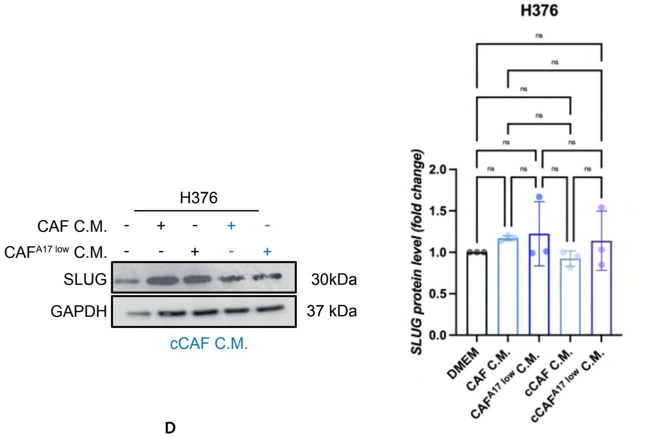


C

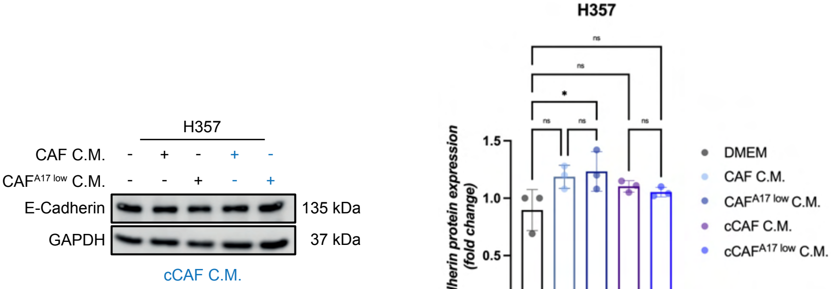


D

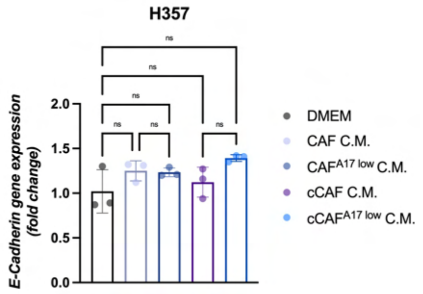


E

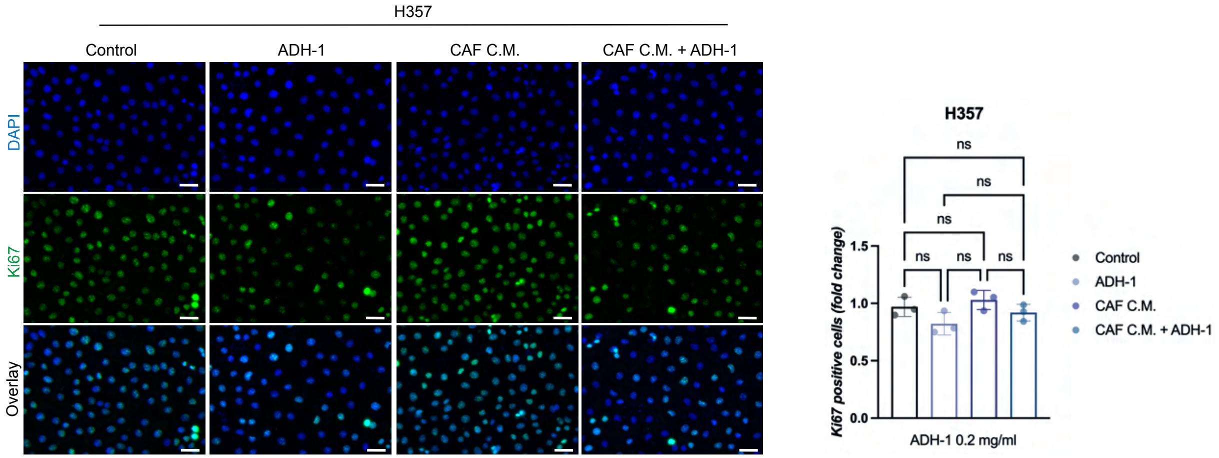


F

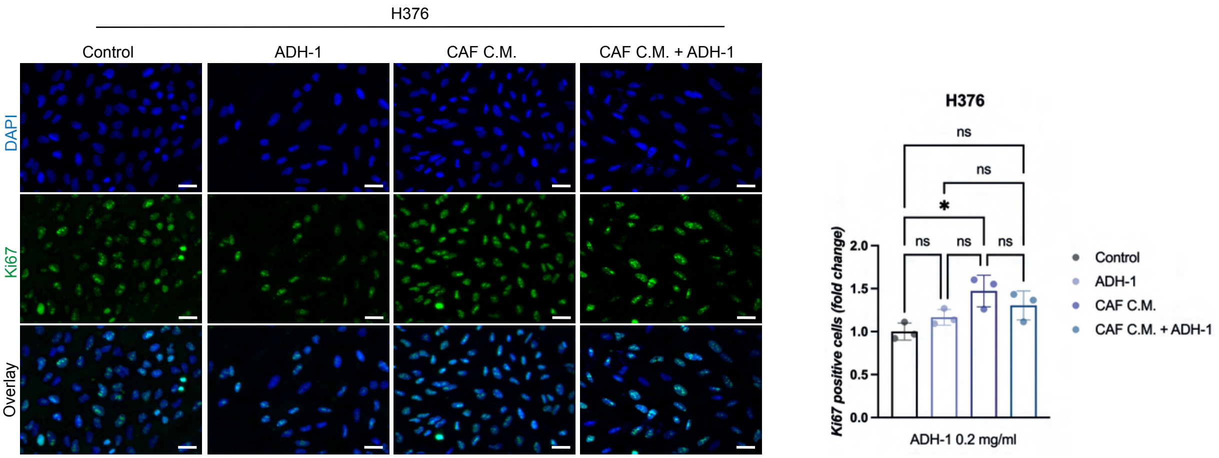

### Supplementary figure 4

#### Supplementary Figure 4

A

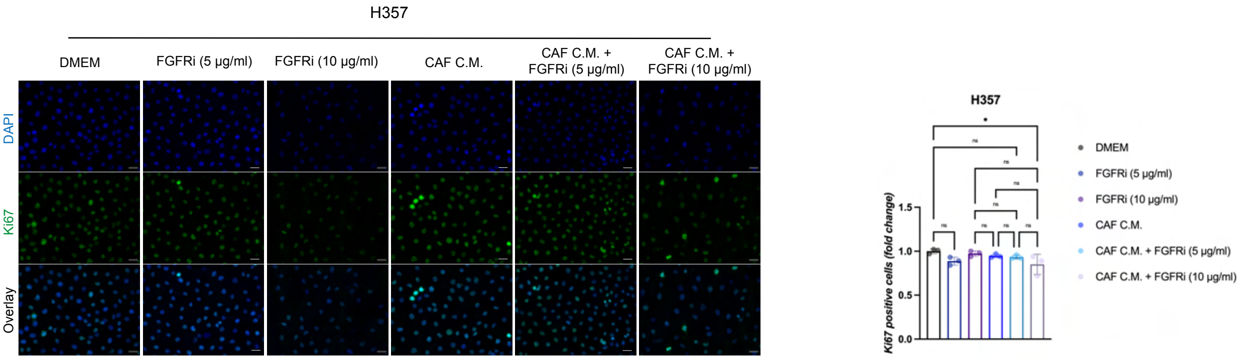


B

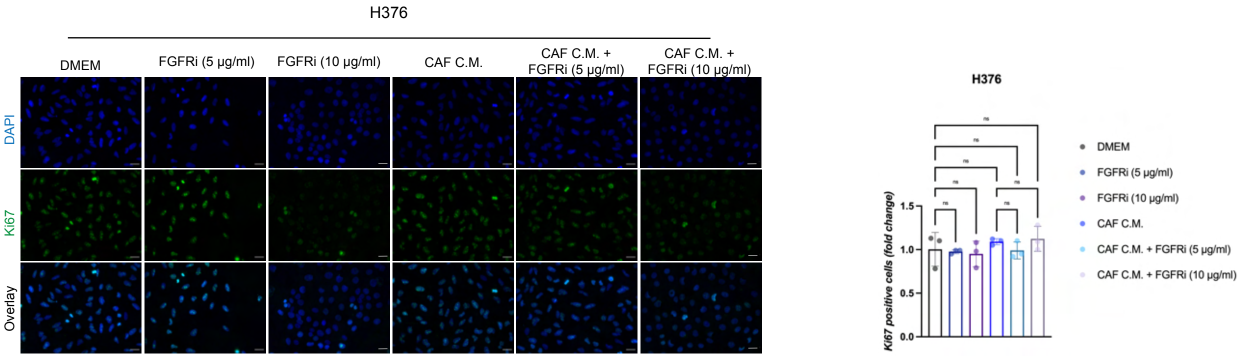


C

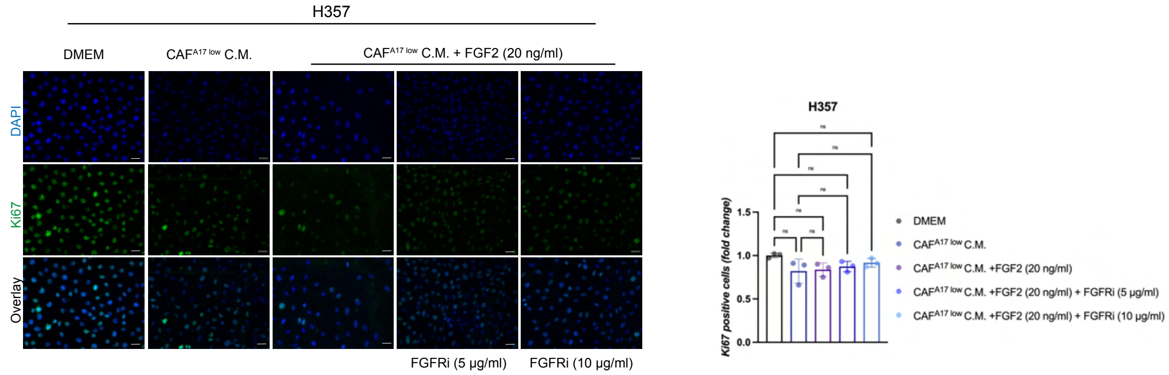


D

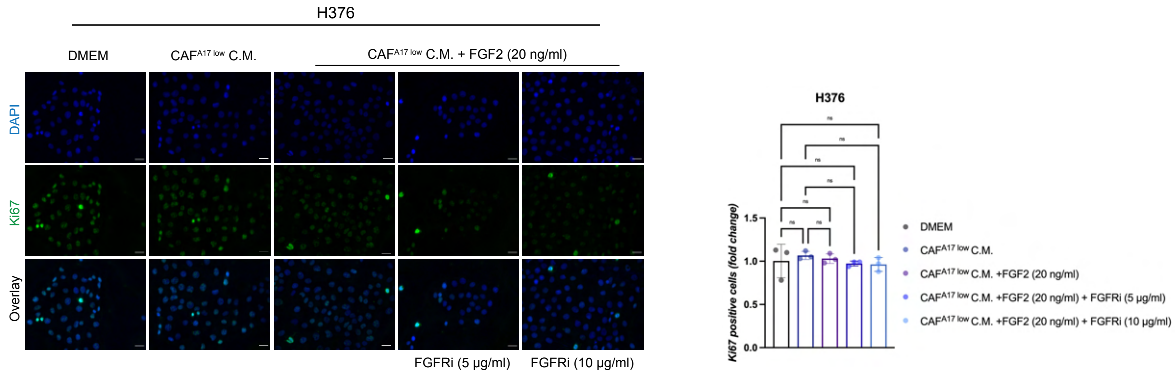
