## Supplementary tables for "Cancer-associated fibroblast ADAM17 mediates a feed-forward loop promoting cancer cell migration"

Supplementary Table 1: STR profile

| | Genotypes | |
|---|---|---|
| **STR locus** | **H357 OSCC Tongue Keratinocyte Cell (Test Sample)** | **H357 (Comparison Sample)** |
| **D5** | 12 \| 14 | 12 \| 14 |
| **D13** | 12 \| 13 | 12 \| 13 |
| **D7** | 11 \| 11 | 11 \| 11 |
| **D16** | 11\| 12 | 11\| 12 |
| **vWA** | 17 \| 19 | 17 \| 19 |
| **Amel** | X \| X | X \| Y |
| **TPOX** | 8 \| 11 | 8 \| 11 |
| **CSF1PO** | 11 \| 13 | 11 \| 13 |
| **THO1** | 6 \| 9 | 6 \| 9 |

H357 matching percentage outcome: 97 %

| Match vs. Mis-match |
|---|
| Match |
| Match |
| Match |
| Match |
| Match |
| Mis-match |
| Match |
| Match |
| Match |

| | Ge |
|---|---|
| **STR locus** | **H376 OSCC, floor of mouth (Test Sample)** |
| **D5** | 10\| 12 |
| **D13** | 8\| 8 |
| **D7** | 10 \| 10 |
| **D16** | 11\| 14 |
| **vWA** | 16 \| 18 |
| **Amel** | X \| X |
| **TPOX** | 8 \| 8 |
| **CSF1PO** | 10 \| 11 |
| **THO1** | 6 \| 9 |

H376 matching percentage outcon

| notypes | |
|---|---|
| H376 (Comparison Sample) | Match vs. Mis-match |
| 12 \| 14 | Mis-match |
| 8 \| 11 | Mis-match |
| 8 \| 10 | Mis-match |
| 11\| 14 | Match |
| 16 \| 18 | Match |
| X \| X | Match |
| 8 \| 8 | Match |
| 10 \| 11 | Match |
| 6 \| 9 | Match |

ne: 93 %

Supplementary Table 1

Supplementary Table 2: Catalogue number and sequence of all siRNAs used

| Gene | Catalogue number |
|---|---|
| **ADAM17** | 4390824, s13718, ThermoFisher Scientific |
| | 4390824, s13720 ThermoFisher Scientific |
| **Negative control** | AM4611, ThermoFisher Scientific |

| Sequence | exon |
|---|---|
| 5'-GTAAAAACGAAAGCGAGTACA-3' | 3 |
| 5'-TTGAACGATTTTGGGATTTCA-3' | 16 |
| Undisclosed (double check) | |

Supplementary Table 3: List of Antibodies used for western blotting

| Taqman primers | |
|---|---|
| **Gene** | **Assay ID and catalogue number** |
| **ADAM17** | Hs01041915_m1, 4331182 |
| **Collagen I** | Hs00164004_m1, 4331182 |
| **aSMA** | Hs05005339_m1, 4351372 |
| **Vimentin** | Hs00958111_m1, 4331182 |
| **Snail** | Hs00195591_m1, 4331182 |
| **Twist** | Hs04989912_s1, 4331182 |
| **Slug** | Hs00161904_m1, 4331182 |
| **N-cadherin** | Hs00983056_m1, 4331182 |
| **FGF2** | Hs00266645_m1, 4331182 |
| **FAP** | Hs00990791_m1, 4331182 |

| Sybr Green primers | | |
|---|---|---|
| **Gene** | | **Sequence** |
| **ADAM17** | FWD | 5’TGAGGGCAGTTAACCAAACC3’ |
| | REV | 5’ATACACCCACACACCCCACT3’ |
| **Collagen I** | FWD | 5’ATGTAGGCCACGCTGTTCTT3’ |
| | REV | 5’AGAGCATGACCGATGGATT3’ |
| **Vimentin** | FWD | 5’AGGTGGACCAGCTAACCAAC3’ |
| | REV | 5’ATTCCACTTTGCGTTCAAGG3’ |
| **aSMA** | FWD | 5’GAAGAAGAGGACAGCACTG3’ |
| | REV | 5’TCCCATTCCCACCATCAA 3’ |
| **U6** | FWD | 5’ CTCGCTTCGGCAGCACA 3’ |
| | REV | 5’AACGTTCACGAATTTGCGT 3’ |

Supplementary Table 4: List of primers used for the RT-qPCR

| Primary antibodies | | | |
|---|---|---|---|
| **Protein** | **Catalogue number** | **Provider** | **Dilution** |
| **ADAM17** | ab2051 | Abcam | 1 to 1000 |
| | ab57484 | | |
| **Collagen I** | 39952S | Cell Signaling Technology | 1 to 1000 |
| **aSMA** | ab7817 | Abcam | 1 to 3000 |
| **FAP** | 66562S | Cell Signaling Technology | 1 to 1000 |
| **GAPDH** | 60004-1-Ig | Proteintech | 1 to 5000 |
| **bactin** | A1978-200UL | Sigma Aldrich | 1 to 10000 |
| **N-cadherin** | 14215S | Cell Signaling Technology | 1 to 1000 |
| **SMAD3** | ab40854 | Abcam | 1 to 2000 |
| **Phospho-SMAD3** | ab52903 | Abcam | |
| **FGF2** | ab208687 | Abcam | 1 to 1000 |
